# Pharmacologic AMPK Activation Extends Lifespan in *C. elegans* and Improves Aspects of Healthspan in Mice

**DOI:** 10.64898/2026.04.21.719899

**Authors:** Özlem Altintas, Marie Knufinke, Charlotte G. Mann, P. Kent Langston, Joshua D. Rabinowitz, Coleen T. Murphy, Collin Y. Ewald, Sarah J. Mitchell, Michael R. MacArthur

**Affiliations:** Department of Health Sciences and Technology, ETH Zurich, Schwerzenbach CH-8603; Department of Pathology, Yale School of Medicine, New Haven CT, 06510 USA; Yale Center for Research on Aging, Yale School of Medicine, New Haven CT, 06510 USA; Department of Chemistry, Princeton University, Princeton NJ 08544 USA; Lewis-Sigler Institute for Integrative Genomics, Princeton University, Princeton NJ 08544 USA; Ludwig Institute for Cancer Research, Princeton University, Princeton NJ 08544 USA; Department of Molecular Biology, Princeton University, Princeton NJ 08544 USA

## Abstract

Dysregulated energy metabolism is a hallmark of aging. Many interventions that extend lifespan converge on the conserved master regulator of energy metabolism, AMP-activated kinase (AMPK), and direct genetic activation of AMPK extends lifespan in multiple species. Here, we test the ability of a specific and potent pharmacologic AMPK activator, MK-8722, to extend lifespan in *C. elegans* and improve healthspan in aged mice. Treatment with MK-8722 from adulthood significantly extended lifespan in an AMPK-dependent manner in both wildtype and Cockayne syndrome model *csb-1* mutant C. elegans, without impairing motility or reproductive capacity. Mice treated with MK-8722 from 18 until 24 months of age had significantly reduced body fat accumulation, blocked age-associated declines in fasting blood glucose and enhanced circadian rhythmicity in respiratory quotient, suggesting an improved overall metabolic state. Hepatic RNA sequencing revealed a decrease in inflammation-related pathways and an increase in sterol metabolic pathways, which was consistent with significantly increased levels of multiple sterol-derived metabolites, including lithocholic acid, a proposed mediator of the benefits of caloric restriction. Our results support pharmacologic AMPK activation as a promising gerotherapeutic strategy.

## Introduction

Metabolic dysfunction is a key feature of aging and includes impaired nutrient sensing, altered mitochondrial function and loss of normal circadian rhythmicity in macronutrient utilization (López-Otín et al., 2013, 2023). Many of the most effective interventions for extending lifespan and healthspan in model organisms broadly affect metabolism, including dietary caloric or amino acid restriction. AMP-activated protein kinase (AMPK) is a central regulator of metabolism. It is activated in response to energy or nutrient stress conditions that increase ATP consumption or interfere with ATP production. In response to decreased cellular energy levels, AMPK is activated through the phosphorylation of its catalytic alpha subunit (Hardie, 2011). In mammals, AMPK exists in 12 different isoforms with tissue- and cell-specific expression patterns (Pinter et al., 2012). The activation of AMPK has widespread effects on metabolism via phosphorylation of downstream target proteins (Hardie, 2014).

In addition to regulating systemic metabolism, AMPK also plays a key role in modulating lifespan. Dietary restriction protocols that increase lifespan and healthspan often drive AMPK activation (Edwards et al., 2010; Palacios et al., 2009) and AMPK activation is required for lifespan extension in multiple worm and fly models of dietary restriction (Greer et al., 2007; Stenesen et al., 2013). Overexpression or direct genetic activation of AMPK via expression of a truncation mutant containing the catalytic domain is sufficient to extend lifespan in worms and flies (Apfeld et al., 2004; Mair et al., 2011; Stenesen et al., 2013; Weir et al., 2017). AMPK is also implicated in some progeroid syndromes, including Cockayne syndrome (CS). Transcriptomic analyses suggest that AMPK signaling is impaired in multiple CS models, including *C. elegans*, mice, and patient-derived tissue samples (Okur et al., 2020). Interventions that indirectly activate AMPK including caloric and methionine restriction also increase lifespan in CS mouse models (Brace et al., 2013; Vermeij et al., 2016).

Considering AMPK’s clear involvement in normal aging, progeroid syndromes and geroprotective interventions, pharmacologic activation of AMPK represents a promising strategy for improving lifespan and healthspan (Burkewitz et al., 2014; Cantó & Auwerx, 2011). Multiple indirect methods of activating AMPK via drugs or supplements, including metformin, thiazolidinediones, berberine and AICAR have shown promise in aging models. However, studies testing direct and specific pharmacologic AMPK activation are lacking. MK-8722 is a potent, highly specific pan-AMPK activator that has robust beneficial effects in preclinical models of metabolic disorders (Myers et al., 2017). In this study, we tested the effects of direct pharmacologic AMPK activation, via MK-8722 treatment, on the lifespan of wild-type and CS model *C. elegans* and on multiple healthspan parameters in wild-type aged mice. We find that MK-8722 extends lifespan in wild-type and CS model *C. elegans* without impairing motility or reproductive capacity. In aged mice, MK-8722 treatment improves multiple healthspan parameters, including circadian rhythmicity of metabolism and liver function. Transcriptomic and metabolomic analysis showed signatures of decreased inflammation and increased sterol-related metabolism, including increased levels of lithocholic acid, a proposed mediator of caloric restriction benefits (Qu, Chen, Wang, Long, et al., 2025). These findings collectively suggest that pharmacologic AMPK activators like MK-8722 may hold promise as gerotherapeutic agents.

## Results

### MK-8722 extends lifespan in wild-type and CS model *csb-1(ok2335)* mutant *C. elegans* in an AMPK-dependent manner

To test the effects of MK-8722 on lifespan, we treated wild type N2 *C. elegans* starting at late L4 stage with concentrations ranging from 10-350 μM and found that 70 μM was the optimal concentration for significantly extending lifespan (**Figure 1A**). Animals were transferred to fresh vehicle or drug plates at days 5 and 10, which was required for maximal efficacy. We also tested 70 μM MK-8722 treatment in AMPK knockout *aak-2* mutant and constitutively active AMPK CA-AAK2 strains. While the *aak-2* mutants were short lived and CA-AAK2 were long lived, as previously reported, 70 μM MK-8722 treatment had no effect in either genotype, confirming on-target effects (**Figure 1A**). We next tested the effects of MK-8722 in a *C. elegans* model of a premature aging syndrome using the *csb-1(ok2335)* mutant, which displays multiple CS-related phenotypes including hypersensitivity to UV, accumulation of dysfunctional mitochondria, and transcriptomic alterations that overlap with CS patient cerebellar samples (Lopes et al 2020; Okur et al., 2020). We treated wild type N2 and *csb-1(ok2335) C. elegans* starting at late L4 stage with 70 μM MK-8722 and observed significant lifespan extension in both genotypes (**Figure 1B**). We also validated that lifespan extension was on-target for AMPK in the CS model by crossing the *aak-2* null and CA-AAK2 alleles into the *csb-1* mutant background and observed similar results as in the N2 background (**Figure S1A**). Many interventions that extend lifespan reduce aspects of fitness in early life. We thus tested whether MK-8722 treatment affected reproductive capacity as a major fitness parameter. We did not observe any effect of MK-8722 treatment on dynamics of reproduction in either selfed (**Figure 1C**) or mated (**Figure S1B**) wild type N2 or *csb-1* animals, and cumulative lifetime brood size was not significantly affected by drug treatment in selfed or mated animals of either genotype (**Figure 1D**).

**Figure 1.**
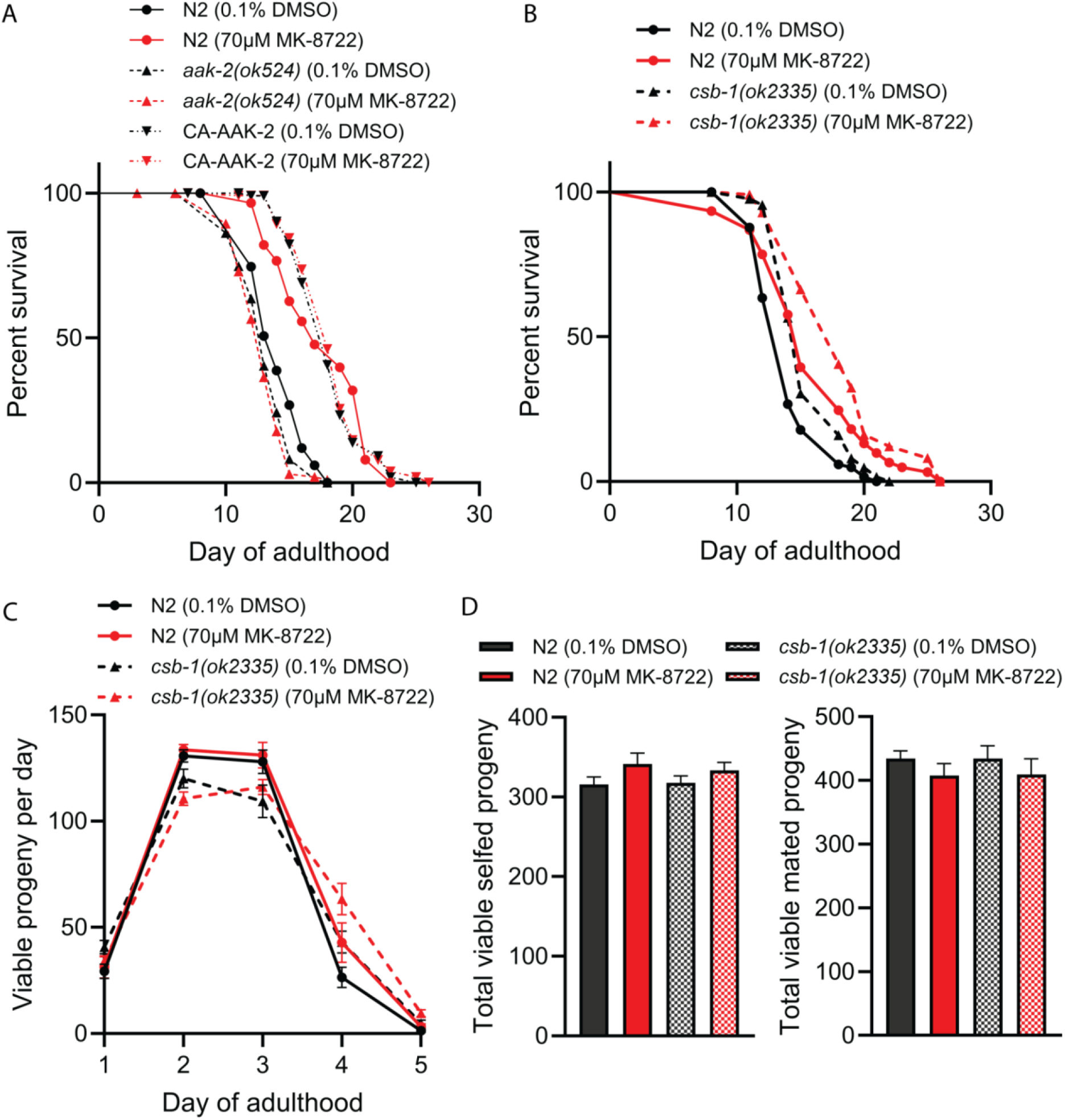
MK-8722 causes AMPK-dependent lifespan extension without affecting reproduction. **A)** Lifespans of wild-type N2, *aak-2* null and CA-AAK-2 (expressing constitutively active AMPK) *C. elegans* treated with vehicle or 70 μM MK-8722. **B)** Lifespans of wild-type and *csb-1* null mutant *C. elegans* treated with vehicle or 70 μM MK-8722. **C)** Total viable progeny per day of self-mating wild-type N2 and *csb-1* null mutant hermaphrodites treated with either vehicle or 70 μM MK-8722. **D)** Total cumulative brood size summed across all days from self-mating (left) or mated (right) *C. elegans*. For lifespans at least 60 worms per condition were used. Detailed statistics and sample sizes are in Supplemental Workbook 1 and methods.

We also validated manually scored survival results with an orthogonal automated method, which uses lifelong imaging on flatbed scanners to assess motility and lifespan. Although we could not refresh drug at days 5 and 10 as in manually scored assays, we still found that 70 μM MK-8722 significantly extended the age at 25% mortality, representing a “squaring of the curve” (**Figure S1C**). We also found a significant extension of lifespan spent displaying high motility, representing an extension of life spent in a healthy state (**Figure S1D**). Detailed statistics for all lifespan experiments are provided in **Supplemental Workbook 1**.

### MK-8722 improves healthspan measures in aging mice

We next tested the effects of MK-8722 on aging phenotypes in mice. We treated 18 month old male and female C57BL/6JRj mice with either control purified diet (Research Diets D12450B, Control) or purified diet supplemented with MK-8722 (MK) at 50 parts per million (ppm) to achieve a daily dose of approximately 4 mg/kg body weight. Mice were sacrificed at 24 months of age after six months of diet treatment. We measured multiple age-related phenotypes including body weight, fat and lean mass, food intake, frailty index, energy expenditure, respiratory exchange ratio (RER) and blood parameters including glucose and liver enzymes.

Mice on MK diet gained significantly less weight over the course of the study and had significantly lower body fat percentage at both 3 and 6 months of treatment (**Figure 2A-B, S2A**). While the overall effect of MK was highly significant, the effects were more pronounced in females than males. Female mice on MK diet consumed approximately 8% less food over the course of the study, while there was no significant difference in food intake between male groups (**Figure S2B**). Consistent with echoMRI body fat measurements, upon sacrifice both male and female MK groups had significantly lower gonadal adipose tissue depot masses (**Figure S2C**). Prior literature has reported increased heart and kidney mass upon genetic or pharmacologic activation of AMPK (Myers et al., 2017; Wilson et al., 2021; Yang et al., 2016). We observed significantly increased heart and kidney masses in MK-treated females, but no differences in MK-treated males (**Figure S2C**). We did not observe any significant time by treatment effects on any echocardiogram parameters, though males on MK did tend to have a lower heart rate at 6 months (**Table S1-2**).

**Figure 2.**
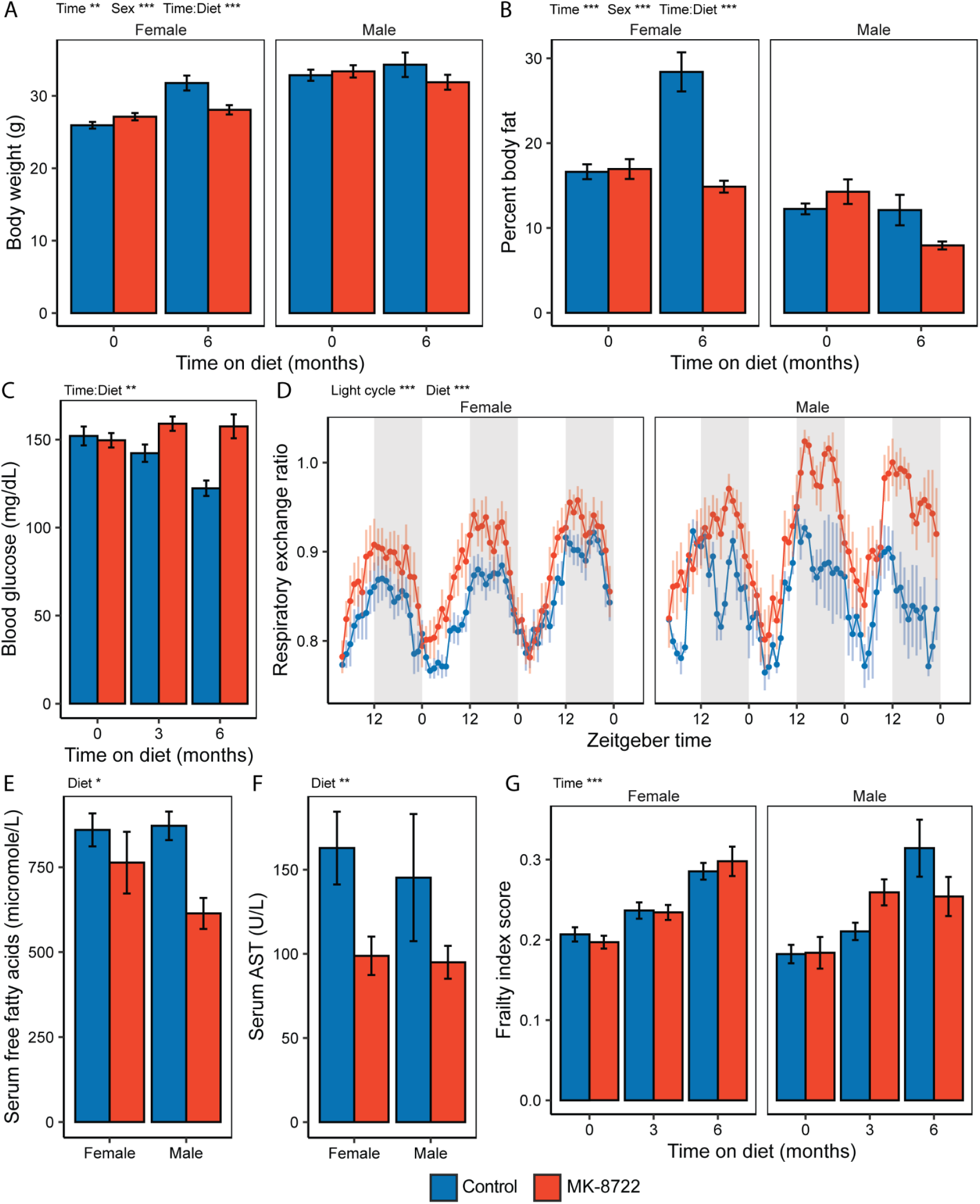
Late-onset MK-8722 treatment improves multiple aging phenotypes in mice. **A)** Body weights at baseline (18 months of age) and after 6 months of treatment (24 months of age) in male and female C57BL/6JRj mice treated with either Control diet or MK-8722 (MK) at 50ppm in the diet. **B)** Percent body fat measured by EchoMRI at baseline and 6 months of treatment. **C)** Fasting blood glucose at baseline, 3 months and 6 months of treatment, males and females combined. **D)** Respiratory exchange ratio over 72 hours of mice treated with Control or MK for 5 months, shaded areas indicate dark cycle. **E)** Serum free fatty acid and **F)** aspartate aminotransferase (AST) levels from samples collected at the time of sacrifice following 6 months of treatment. **G)** Frailty index scores at baseline, 3 months and 6 months of treatment. For panels with multiple timepoints asterisks indicate term significance from ANOVA including sex, time, diet and time-by-diet interaction. For panels with one timepoint asterisks indicate term significance from ANOVA including sex and diet. *, **, *** indicate P<0.05, 0.01, 0.001 respectively. Group N was 7-20, as detailed in methods.

Reduced fasting blood glucose is a feature of aging in mice (Palliyaguru et al., 2021). Treatment with MK completely blocked the age-associated suppression in fasting blood glucose observed in Control mice (**Figure 2C**). Treatment with MK also significantly increased the respiratory exchange ratio at 5 months of treatment, particularly during the dark cycle, suggesting improved maintenance of circadian rhythmicity in metabolism (**Figure 2D, S2D**). Per-mouse total energy expenditure and locomotor activity were not significantly affected by MK treatment. Serum free fatty levels were significantly lower in MK-treated mice after 6 months of treatment (**Figure 2E**). Liver damage markers were also improved by MK treatment, with mice on MK showing significantly reduced serum AST and ALT (**Figure 2F, S2E**). Serum urea and creatinine were not significantly different between treatment groups (**Figure S2F-G**), suggesting minimal impairment of renal function. Despite improvements in multiple metabolic aging phenotypes, frailty as measured by the 31-parameter mouse clinical frailty index (Whitehead et al., 2014a) was not significantly different between treatment groups at 3 or 6 months of treatment (**Figure 2G**).

### MK-8722 drives geroprotective shifts in the metabolome of aging mice

Because we identified multiple metabolic phenotypes altered by MK treatment in aging mice, we next performed metabolomic analysis of liver and kidney tissue collected at the time of sacrifice from Control and MK treated males and females. In regression models accounting for sex and treatment, 27 metabolites were significantly altered in the liver and 224 in the kidney by MK treatment after FDR correction (**Figure S3A-B**). In the kidney, many lipid-related compounds were significantly reduced by MK treatment, including multiple saturated fatty acids (C15:0, C16:0, C17:0), two prostaglandins and an acyl-taurine derivative (**Figure 3A, S3C**). Multiple energy-yielding compounds significantly increased in both male and female, including ATP, creatine phosphate and hexose-phosphate (**Figure 3A, S3D**).

**Figure 3.**
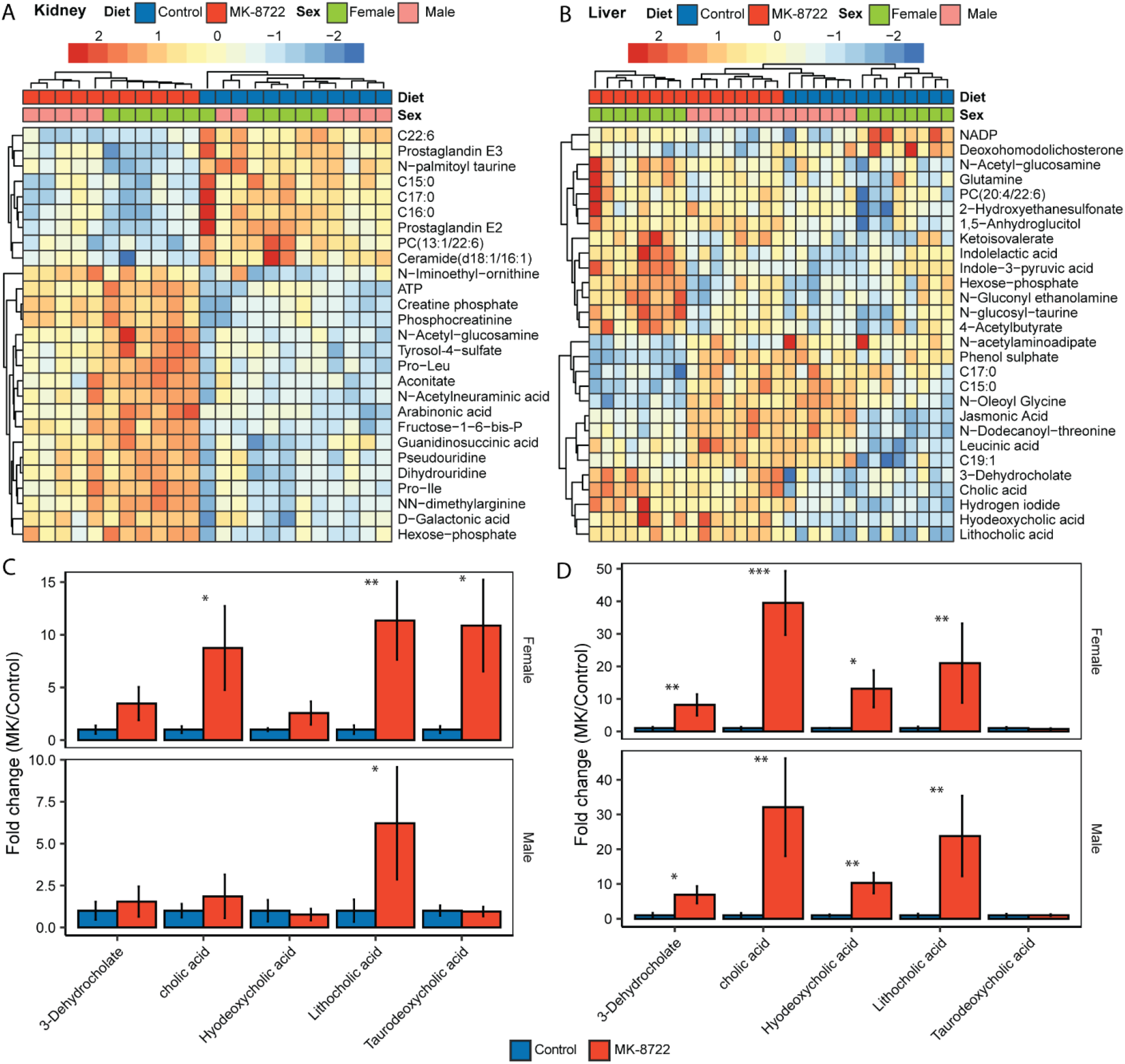
Late-onset MK-8722 drives geroprotective shifts in the metabolome. **A)** Heatmap showing metabolites significantly altered in the kidney at FDR-corrected P<0.001 in 24-month old male and female mice treated with control or MK-8722 for 6 months prior to sacrifice. **B)** Heatmap showing metabolites significantly altered in the liver at FDR-corrected P<0.05. **C)** Fold changes (MK/control) for sterol-derived metabolites in the kidney and **D)** liver. Asterisks *, **, *** indicate P<0.05, 0.01, 0.001 respectively from a non-parametric t-test comparing MK to Control within sex.

The liver showed similar trends for lipid-derived and energy-yielding compounds, but with greater sexual dimorphism (**Figure 3B, S3C-D**). We also noted significant and robust increases in multiple bile acid species including cholic acid, hyodeoxycholic acid, 3-dehydrocholic acid, taurodeoxycholic acid and lithocholic acid in both liver and kidney of male and female mice treated with MK (**Figure 3C-D**).

Many of these sterol-derived metabolites are depleted with aging and display a variety of bioactivities, with lithocholic acid recently noted as a potent geroprotective compound in multiple species (Qu, Chen, Wang, Long, et al., 2025; Qu, Chen, Wang, Wang, et al., 2025).

### MK-8722 effects on the hepatic transcriptome mirror metabolomic responses

We next performed RNA sequencing on liver tissue to better understand potentially regulatory mechanisms underlying the metabolomic responses we observed. We performed sequencing in females because they tended to show larger effect-size responses in metabolomic data. A set of 23 genes were significantly differentially expressed after false detection rate (FDR) correction (**Figure 4A**). Interestingly, among the significant hits, multiple genes involved in sterol metabolism were significantly increased including Cyp1a2, Cyp2c39, Sult1c2, Ugt2a3 and Hsd2b3 (**Figure 4A**). Gene set enrichment analysis revealed that nearly all of the 10 top increasing pathways were involved in sterol or isoprene metabolic pathways (**Figure 4B**). The most decreased pathways were primarily involved in interferon-related inflammatory signaling (**Figure 4B**). The signature relating to enrichment of sterol metabolic pathways was very robust, as seen in the top related pathway entry, Steroid Biosynthetic Process (GO:0006694, **Figure 4C**). While not all genes in this pathway were significantly increased after FDR correction, the trend to increase was robust with 19 genes showing nominally significant increases (**Figure 4D**). A similar case was observed for genes related to interferon-beta signaling (**Figure S4A**), where 15 genes decreased with nominally significant p-values (**Figure S4B**).

**Figure 4.**
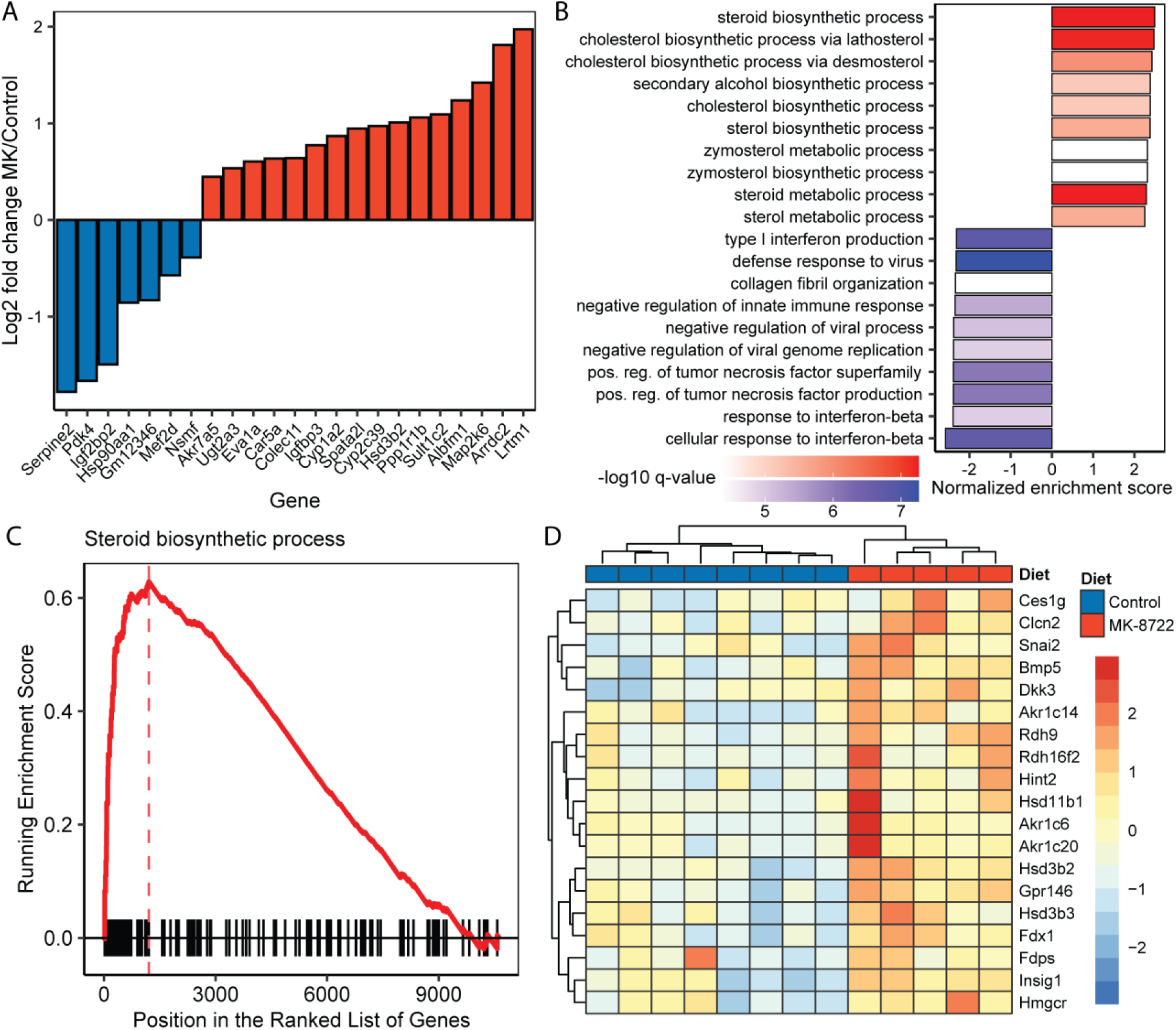
Late-onset MK-8722 drives sterol and inflammation-related transcriptional signatures in the liver. **A)** Log2 fold changes for all genes that were significantly differentially expressed in 24 month old female mice following 6 months of MK treatment. **B)** Top 10 GO pathways with the largest positive and negative normalized enrichment scores from gene set enrichment analysis (GSEA). **C)** GSEA plot for the GO pathway “Steroid Biosynthetic Process”. **D)** Heatmap of all nominally significantly expressed genes in the GO pathway “Steroid Biosynthetic Process”.

## Discussion

In this study, we tested the lifespan and healthspan effects of MK-8722, a specific and potent pan-AMPK activator, in normal aging using N2 *C. elegans* and wild-type mice, and in the *C. elegans csb-1(ok2335)* model of the progeroid disease Cockayne Syndrome. We found that life-long treatment with MK-8722 at 70 μM significantly extends lifespan by approximately 13% in both genotypes, and that this extension is AMPK-dependent. The period of life during which animals showed vigorous motility was also extended by MK-8722 treatment, suggesting increased healthspan in addition to lifespan. We also examined whether MK-8722-mediated lifespan extension adversely affects reproduction, as a potential trade-off between lifespan extension and reproductive fitness seen in some long-lived mutants (Gems et al., 1998; Zhang et al., 2022). We observed no significant changes in daily or total brood size with either selfing or mating following MK-8722 treatment. We also observed enhanced motility in animals treated with MK-8722. These data suggest that the lifespan extension conferred by MK-8722 does not incur an obvious fitness cost, as seen in some other models of lifespan extension.

In our hands, *csb-1(ok2335)* mutant *C. elegans* did not live shorter than wild-type. While some previous work has reported shorter lifespan in the *csb-1(ok2335)* strain (Okur et al., 2020), other work has reported no difference or even increased lifespan in *csb-1(ok2335)* and other DNA repair-deficient *C. elegans* strains (Lans et al., 2013; Lopes et al., 2020). We speculate that these discrepancies could be driven by different genetic backgrounds, as null alleles are often crossed onto lab-specific wild-type strains, or lab-specific differences in environmental or maintenance conditions. Literature does consistently show UV sensitivity in *csb-1(ok2335)* mutants and future work could test the effects of MK-8722 on UV-induced phenotypes.

In mice, MK-8722 protected against multiple age-associated metabolic phenotypes. Mice on MK-8722 accumulated less body fat, sustained their blood glucose levels and had enhanced circadian rhythmicity in respiratory quotient. As we did not observe an increase in total energy expenditure, we conclude that reduced body mass and adiposity is likely driven by mild but sustained suppression in food intake. This food intake suppression may be caused by peripheral effects on insulin sensitivity. Central effects also cannot be ruled out; the brain penetrance of MK-8722 has not been reported but preliminary results from another group show that orally-delivered MK-8722 does increase ACC1 phosphorylation in mouse brain.

While MK-8722 showed striking anti-diabetic effects in preclinical models, on-target cardiac and renal hypertrophy limited clinical development. We also observed significant increases in heart and kidney weight in MK-8722 treated female but not male mice. However, we did not observe any significant time-by-treatment effects in any echocardiogram parameters, and also did not observe any significant changes in serum urea or creatinine levels. While these hypertrophic effects do need to be carefully assessed, a lack of associated functional effects is encouraging. Preclinical work suggests that increased heart weight is reversible and partially driven by glycogen accumulation. The MK-8722 induced cardiac phenotype is reminiscent of athletic heart syndrome, which is broadly considered non-pathologic and is not associated with reduced life expectancy.

Molecular profiling revealed significant changes in hepatic and renal metabolomes with MK-8722 treatment. Many energy-yielding metabolites including ATP, hexose-phosphate and creatine-phosphate were significantly increased in the liver by MK-8722. Many free fatty acids and acylated metabolites were also decreased in tissues from MK-8722 treated mice. Overall, these changes are suggestive of an improved metabolic state in MK-8722 treated animals. We also observed striking increases in many sterol-derived bile acids including lithocholic acid. We previously reported that circulating lithocholic acid is depleted in aging mice (Jankowski et al., 2025). Recent work has also shown that lithocholic acid is induced by caloric restriction and that treating aged mice with lithocholic acid phenocopies many of the geroprotective effects of caloric restriction and extends lifespan.

Mechanistically, lithocholic acid activates AMPK via TULP3 and extends lifespan in *C. elegans* and *D. melanogaster* in an AMPK-dependent manner. Our results suggest that the relationship between lithocholic acid and AMPK may be bidirectional; in addition to lithocholic acid activating AMPK, AMPK activation may increase lithocholic acid levels. We also observed increases in other sterol-derived bile acids, and thus hypothesize that the increase in lithocholic acid by AMPK activation is likely not specific, but rather downstream of general modification of bile acid synthetic pathways. This is supported by RNA sequencing results showing consistently increased expression of sterol-related genes in response to MK-8722.

While our data show strong geroprotective activities of MK-8722, some limitations should be noted. First, we did not measure lifespan in mice. While we speculate, based on functional measurements, that the cardiac and renal hypertrophy induced by MK-8722 may not be pathologic, we cannot definitively conclude whether these hypertrophies may ultimately negatively affect lifespan.

Second, we did not test the effects of MK-8722 on UV-induced phenotypes in *csb-1* mutant *C. elegans*. UV-irradiation of *csb-1* mutants produces a phenotype that more closely resembles human Cockayne Syndrome, including shortened lifespan and ataxia. Future studies in irradiated *csb-1* mutants or in Cockayne Syndrome mouse models will be useful in determining whether MK-8722 may hold therapeutic potential in Cockayne Syndrome.

## Supporting information

Supplemental Workbook 1

## Acknowledgements

We thank James Vath, Robert Myers, Huseyin Mehmet, Min-woo Kim and Jaret Malloy for their helpful input throughout the project. This work was supported by the National Institute on Aging (P01AG055369 to S.J.M.), and ETH Zurich Doc. Mobility Fellowship to Ö.A.

## Supplemental Figures

**Supplementary Figure 1.**
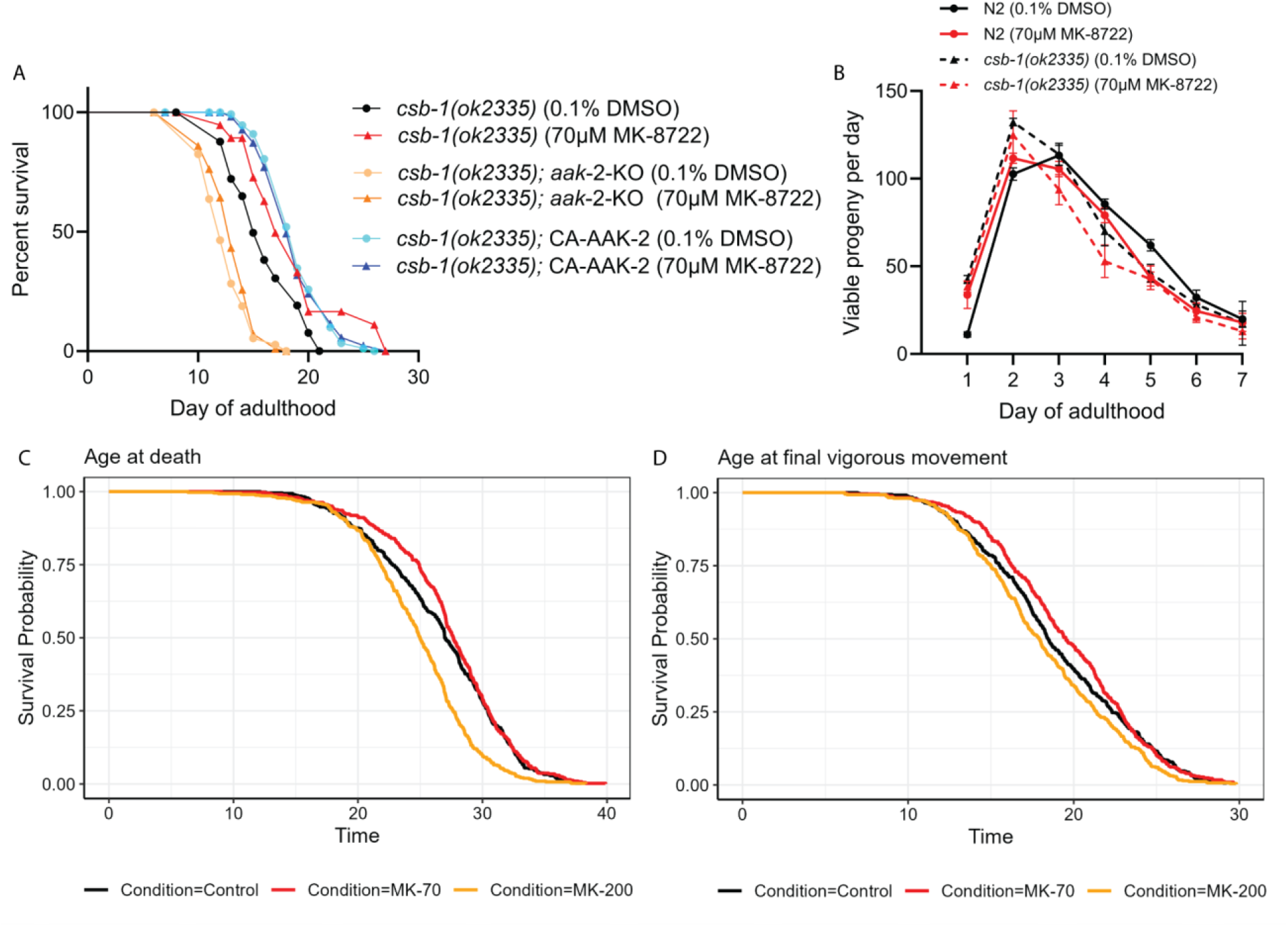
**A)** Total viable progeny per day from day 1 until day 7 of adulthood of wild-type and *csb-1(ok2335)* hermaphrodites mated to males of the same genotype and treated with vehicle or 70 μM MK-8722. **B)** Lifespans of *csb-1(ok2335)* mutants, *csb-1(ok2335)* mutants crossed with *aak-2* null mutants and *csb-1(ok2335)* mutants expressing constitutively active AAK-2 and treated with vehicle or 70 μM MK-8722. **C)** Survival curves for wild-type N2 *C. elegans* treated with vehicle control or MK-8722 at 70 μM or 200 μM with analysis and quantification performed using an automated image-based Worm Lifespan Machine protocol. **D)** Quantitation of age of final vigorous movement from the experiment described in **C**. For manual lifespans at least 60 worms per condition were used. Detailed statistics and sample sizes are in Supplemental Workbook 1 and methods.

**Supplementary Figure 2.**
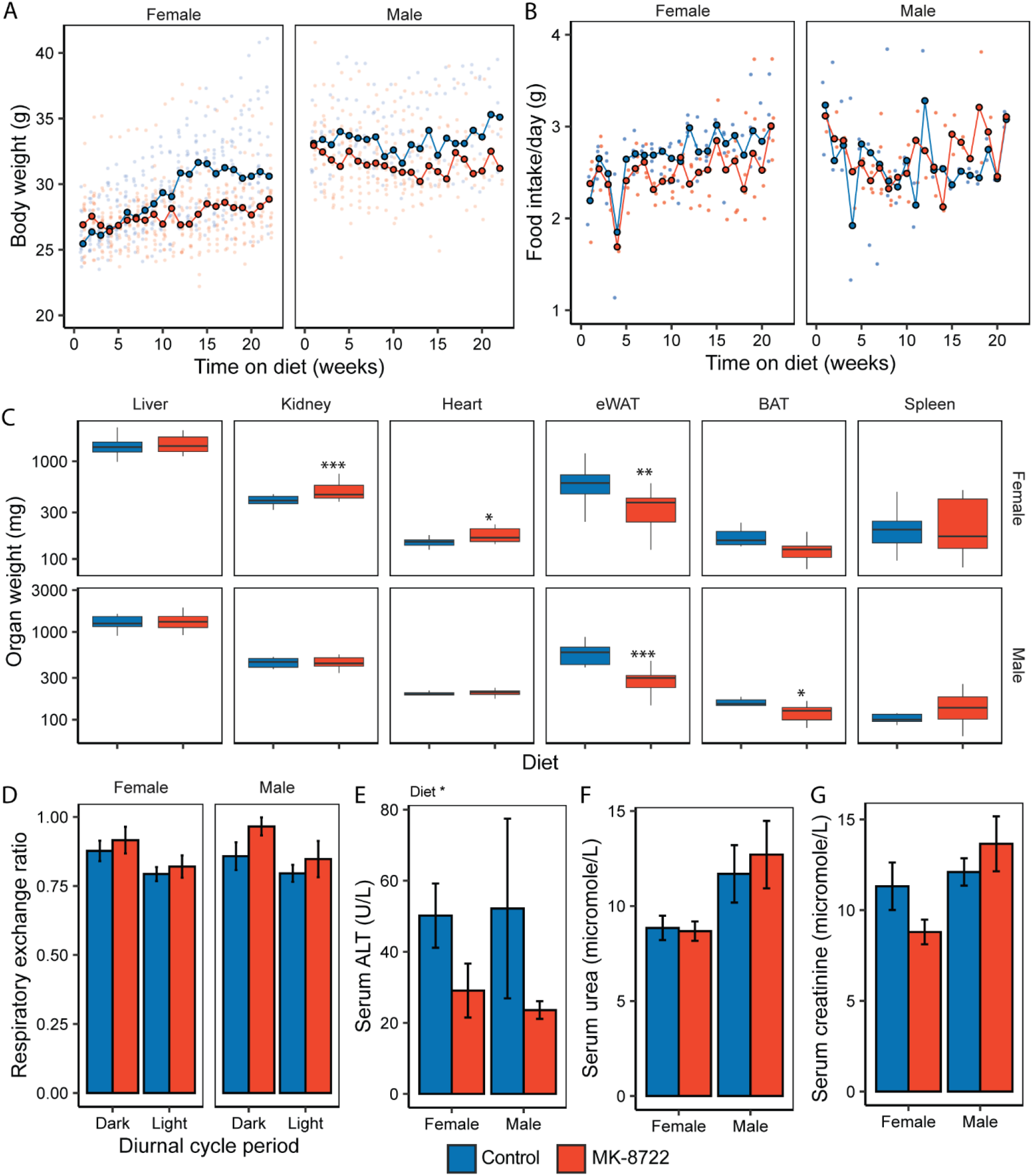
**A)** Body weight trajectories of control and MK-treated female and male mice over the six-month study period. **B)** Food intake of control and MK-treated female (left) and male (right) mice over the six-month study period. **C)** Organ weight of control and MK-treated female and male mice following sacrifice at the end of the six-month study period. **D)** Respiratory exchange ratio of control and MK-treated female and male mice after approximately 5 months of treatment. Values are the per-mouse median across three days of measurement for the 12-hour dark and 12-hour light cycles. **E)** Serum alanine aminotransferase (ALT), **F)** urea, and **G)** creatinine concentrations from serum collected at the time of sacrifice. For **C)** asterisks indicate significance from non-parametric t-test comparing MK to Control within sex. For E-G asterisks indicate term significance from ANOVA including sex and diet. *, **, *** indicate P<0.05, 0.01, 0.001 respectively. Group N was 7-20, as detailed in methods.

**Supplementary Figure 3.**
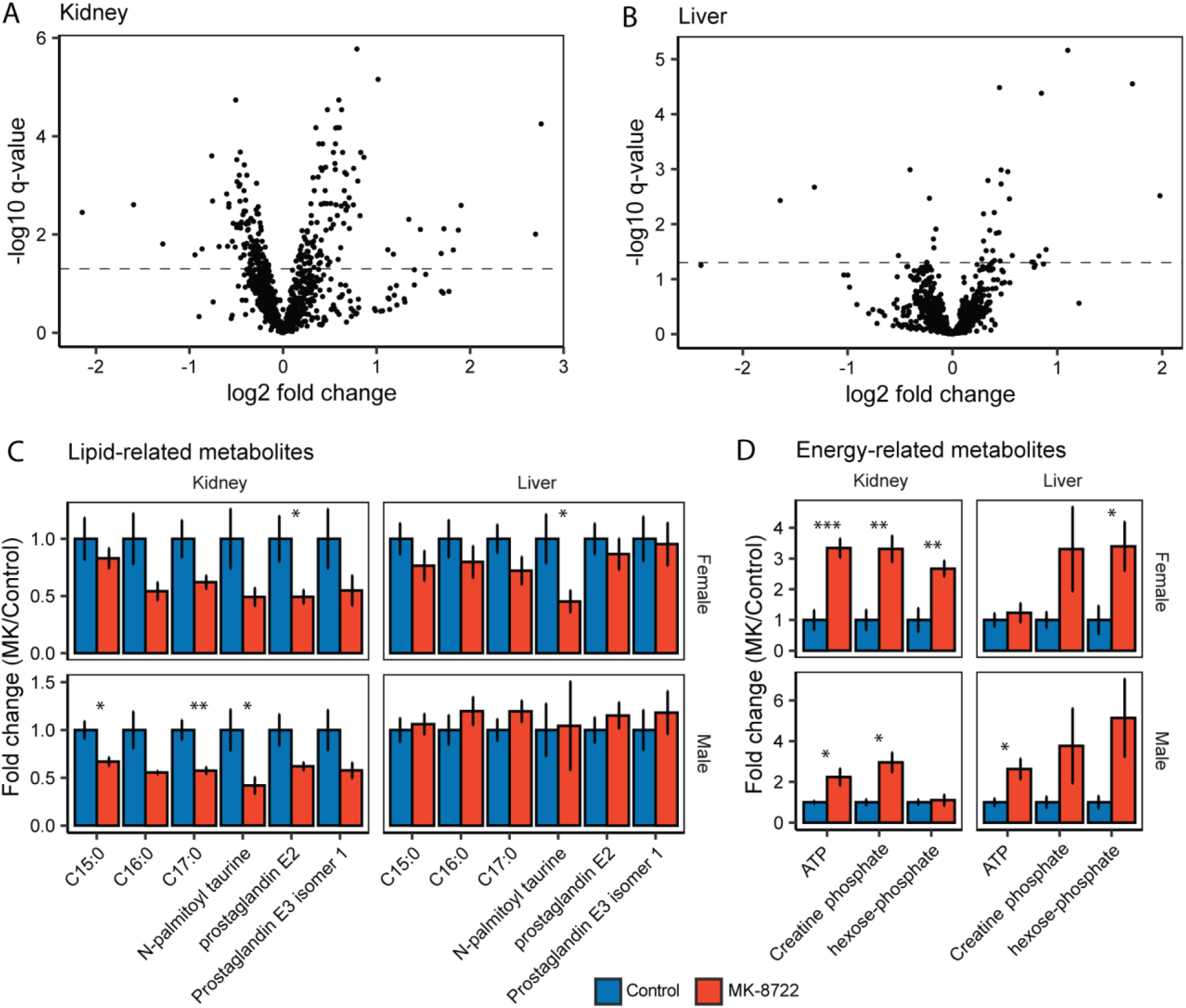
**A)** Volcano plot showing log2 fold change (MK/Control) and −log10 FDR-corrected p-values for polar metabolites measured in the kidney and **B)** liver of male and female mice treated with MK or Control for 6 months from 18-24 months of age. **C)** Fold changes vs Control treated mice of lipid-related and **D)** energy production-related metabolites that showed a significant diet-effect (MK vs Control) in statistical models assessing effects of diet, sex and MK-by-sex interaction in kidney and liver metabolomics data. Asterisks *, **, *** indicate P<0.05, 0.01, 0.001 respectively from a non-parametric t-test comparing MK to Control within sex.

**Supplementary Figure 4.**
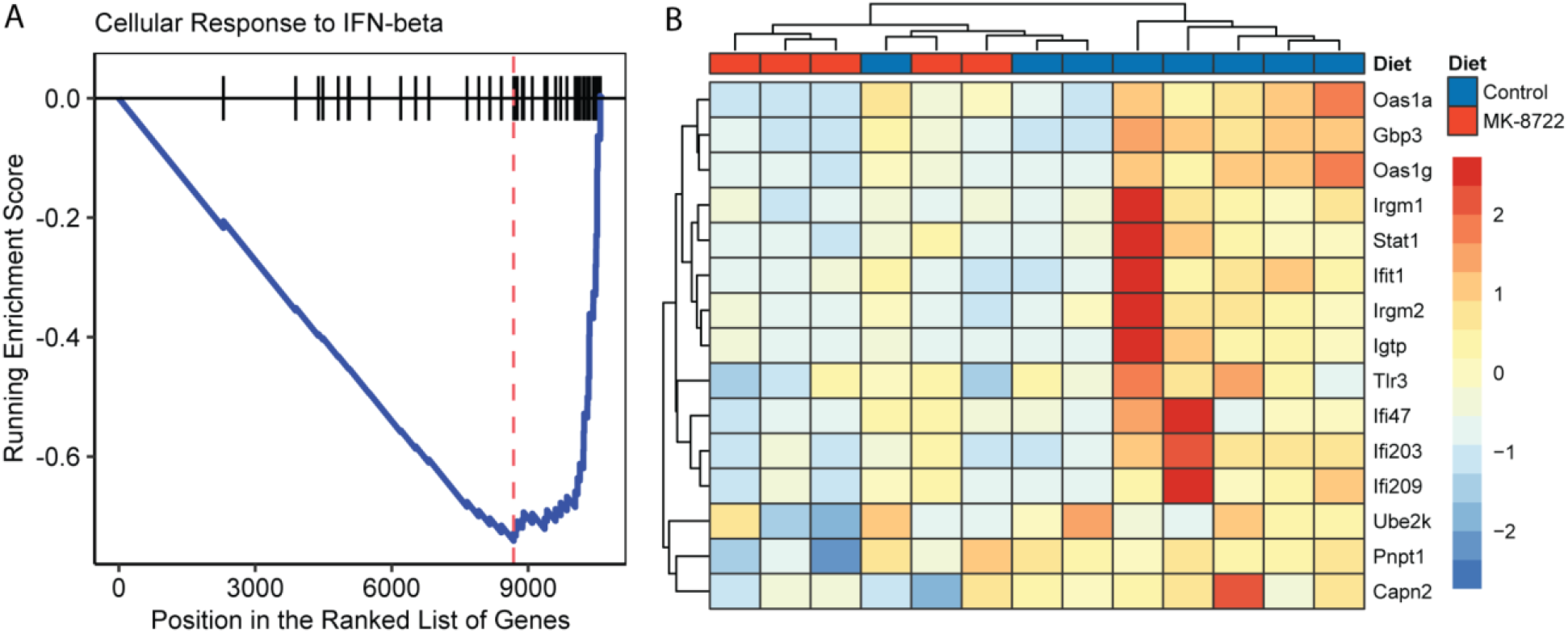
**A)** GSEA plot for the GO pathway “Cellular Response to Interferon-Beta”. **B)** Heatmap of all nominally significantly expressed genes in the GO pathway “Cellular Response to Interferon-Beta”.

**Table S1.**
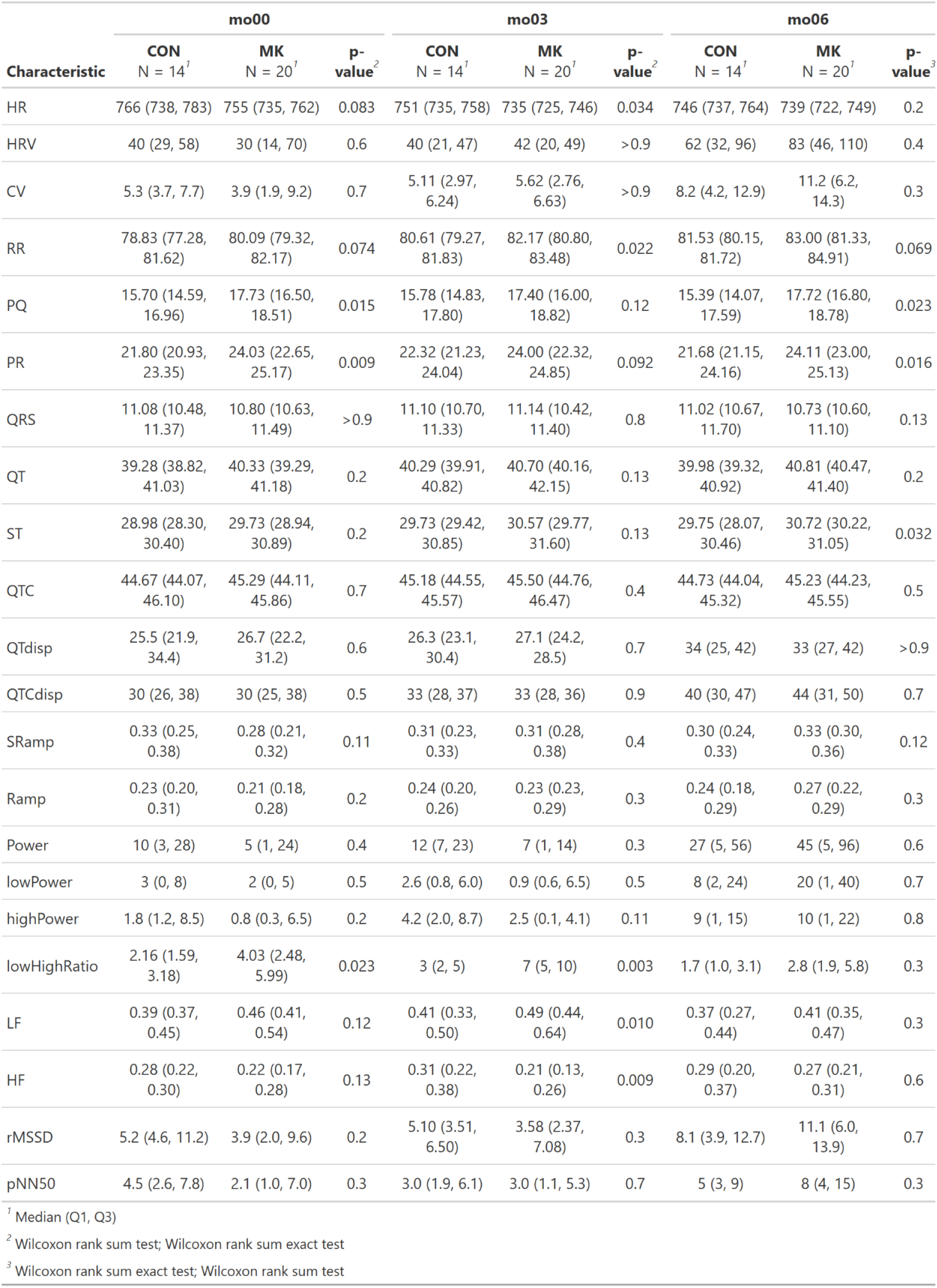
Electrocardiogram parameters in female mice.

**Table S2.** Electrocardiogram parameters in male mice.

| Characteristic | mo00 |  |  | mo03 |  |  | mo06 |  |  |
| --- | --- | --- | --- | --- | --- | --- | --- | --- | --- |
|  | CON<br>N = 7<sup>1</sup> | MK<br>N = 14<sup>1</sup> | p-value<sup>2</sup> | CON<br>N = 7<sup>1</sup> | MK<br>N = 14<sup>1</sup> | p-value<sup>2</sup> | CON<br>N = 7<sup>1</sup> | MK<br>N = 14<sup>1</sup> | p-value<sup>3</sup> |
| HR | 755 (745, 784) | 757 (744, 774) | >0.9 | 751 (738, 772) | 741 (718, 750) | 0.14 | 758 (747, 774) | 744 (740, 751) | 0.073 |
| HRV | 49 (30, 83) | 77 (52, 85) | 0.3 | 36 (19, 64) | 44 (21, 87) | 0.6 | 99 (68, 100) | 87 (42, 136) | 0.7 |
| CV | 6.6 (4.1, 11.0) | 10.7 (6.8, 11.3) | 0.3 | 4.8 (2.6, 8.3) | 6.0 (2.9, 11.9) | 0.5 | 13 (9, 13) | 11 (6, 18) | 0.7 |
| RR | 80.07 (77.60, 80.82) | 79.77 (78.50, 81.83) | 0.6 | 80.40 (78.33, 81.64) | 81.56 (80.60, 84.56) | 0.10 | 81.17 (78.62, 82.34) | 81.78 (80.48, 84.20) | 0.3 |
| PQ | 17.10 (15.13, 19.80) | 16.69 (14.99, 18.03) | 0.5 | 16.59 (15.73, 17.78) | 17.20 (16.26, 18.70) | 0.4 | 16.02 (14.24, 18.20) | 17.22 (15.58, 17.95) | 0.9 |
| PR | 22.97 (21.80, 25.70) | 22.54 (21.63, 23.78) | 0.4 | 22.29 (22.08, 24.28) | 23.23 (22.07, 24.36) | 0.6 | 22.65 (20.82, 23.62) | 23.13 (21.80, 24.23) | 0.6 |
| QRS | 10.45 (10.13, 11.14) | 10.75 (10.25, 11.14) | 0.6 | 10.88 (10.07, 11.05) | 10.13 (9.80, 10.62) | 0.2 | 10.79 (10.48, 10.97) | 11.06 (10.67, 11.26) | 0.4 |
| QT | 39.56 (39.30, 39.98) | 40.31 (39.27, 41.40) | 0.5 | 40.30 (39.00, 40.85) | 39.68 (39.48, 41.45) | >0.9 | 39.78 (39.68, 40.85) | 40.55 (39.90, 41.75) | 0.4 |
| ST | 29.80 (28.94, 30.03) | 29.66 (28.81, 31.20) | 0.8 | 29.55 (29.44, 30.65) | 30.38 (29.95, 31.33) | 0.3 | 29.73 (29.33, 30.24) | 30.15 (29.43, 30.40) | 0.4 |
| QTC | 44.53 (44.10, 45.26) | 45.08 (44.34, 45.25) | 0.6 | 45.04 (43.88, 45.30) | 44.28 (43.56, 45.41) | 0.8 | 44.80 (44.30, 45.38) | 44.94 (44.75, 46.47) | 0.7 |
| QTdisp | 28 (23, 31) | 36 (33, 45) | 0.081 | 29 (26, 37) | 32 (29, 35) | 0.7 | 45 (41, 50) | 45 (27, 56) | 0.9 |
| QTCdisp | 35 (30, 42) | 43 (39, 52) | 0.11 | 34 (31, 44) | 36 (32, 46) | 0.8 | 54 (48, 63) | 52 (33, 64) | 0.7 |
| SRamp | 0.31 (0.23, 0.35) | 0.29 (0.19, 0.32) | 0.5 | 0.26 (0.19, 0.29) | 0.29 (0.18, 0.36) | 0.5 | 0.26 (0.25, 0.27) | 0.19 (0.17, 0.22) | 0.3 |
| Ramp | 0.21 (0.17, 0.31) | 0.20 (0.16, 0.27) | 0.3 | 0.22 (0.18, 0.24) | 0.20 (0.17, 0.25) | >0.9 | 0.22 (0.20, 0.23) | 0.18 (0.15, 0.23) | 0.3 |
| Power | 9 (5, 13) | 26 (22, 49) | 0.011 | 3 (1, 27) | 11 (3, 24) | 0.8 | 72 (58, 82) | 94 (9, 157) | 0.8 |
| lowPower | 2 (1, 3) | 10 (3, 13) | 0.046 | 2.0 (0.2, 18.3) | 3.7 (0.8, 8.3) | >0.9 | 24 (17, 31) | 38 (3, 71) | 0.6 |
| highPower | 2 (1, 4) | 11 (5, 16) | 0.037 | 1.3 (0.3, 5.2) | 3.6 (1.1, 8.3) | 0.4 | 15 (11, 31) | 22 (3, 35) | 0.9 |
| lowHighRatio | 1.89 (0.45, 2.55) | 1.67 (1.17, 2.51) | 0.7 | 2.64 (2.10, 4.70) | 1.63 (0.84, 3.33) | 0.12 | 1.50 (0.81, 2.22) | 1.98 (1.67, 2.67) | 0.15 |
| LF | 0.37 (0.17, 0.42) | 0.32 (0.26, 0.38) | >0.9 | 0.41 (0.33, 0.48) | 0.34 (0.23, 0.44) | 0.3 | 0.30 (0.23, 0.40) | 0.36 (0.33, 0.46) | 0.15 |
| HF | 0.35 (0.28, 0.40) | 0.31 (0.28, 0.38) | 0.8 | 0.30 (0.22, 0.36) | 0.31 (0.22, 0.39) | 0.7 | 0.33 (0.27, 0.38) | 0.29 (0.27, 0.32) | 0.3 |
| rMSSD | 6.6 (5.2, 10.6) | 10.6 (7.6, 12.6) | 0.11 | 5.5 (2.7, 8.5) | 6.4 (3.3, 11.4) | 0.7 | 14 (12, 18) | 17 (4, 24) | >0.9 |
| pNN50 | 5 (2, 7) | 7 (5, 10) | 0.3 | 3.0 (1.0, 6.4) | 2.8 (1.1, 6.9) | >0.9 | 17 (6, 26) | 14 (4, 18) | 0.6 |

<sup>1</sup> Median (Q1, Q3)
<sup>2</sup> Wilcoxon rank sum test; Wilcoxon rank sum exact test
<sup>3</sup> Wilcoxon rank sum exact test; Wilcoxon rank sum test

## Materials and Methods

### C. elegans strains

Some of the *C. elegans* strains were provided by the Caenorhabditis Genetics Center (CGC), which is funded by NIH Office of Research Infrastructure Programs (P40 OD010440). The strains used in this study are as follows: wild-type Bristol strain N2, *csb-1(ok2335) X*, RB754 *aak-2(ok524)* X, CA-*aak-2, csb-1(ok2335)*;CA-*aak-2, aak-2*-KO.

### Preparation of nematode growth media (NGM) plates for maintaining *C. elegans*

NGM agar plates were prepared under sterile conditions. 100 mm sterile petri dishes were seeded with 1 ml OP50 (Escherichia coli) culture, 60 mm plates were seeded with 300 μl OP50. The plates were left on the bench to air-dry. Lifespan assays were conducted with 5 μM 5-fluoro-2′-deoxyuridine (FUDR) starting from late L4 stage to prevent eggs from hatching.

### Handling of *C. elegans*

*C. elegans* were kept at 20 °C on nematode growth medium (NGM) agar plates seeded with fresh OP50 as the primary bacterial food source following standard laboratory culture conditions.

### Genotyping primers

The *csb-1(ok2335)* allele was backcrossed into the lab’s wild-type Bristol N2 background. The presence of the *csb-1* null allele was confirmed using the following genotyping primers.

*csb-1(ok2335)*

F1 - 5′ ATCTTGATTTAATCAAATTCAAATTACCATCC 3′

F2 - 5′ TGAGTAGGTGGTGAGGATGATG 3′

R - 5′ TGACAAATTATCAGGTTCTGGTATTTCAC 3′

### Synchronization of *C. elegans* by bleaching

*C. elegans* were grown into gravid adulthood before bleaching. *C. elegans* were washed with M9 buffer, collected in a 15mL Falcon tube, and centrifuged at 1,800 × g for 1 minute. After two additional washes, 3mL bleach solution was added and vortexed every 30 seconds for 2-3 minutes. Animals were checked under a light microscope until bodies were dissolved, exposing the eggs. Bleaching was stopped by adding M9 buffer to a final volume of 15 mL. Eggs were washed with M9 at least 4 times until the bleach odor was no longer present. The washed eggs were transferred onto agar plates for hatching.

### Manual lifespan assays

5-fluoro-2’-deoxyuridine (FUDR) was used to prevent progeny from hatching. Synchronized L4 *C. elegans* were transferred onto freshly prepared 5 μM FUDR-treated NGM plates seeded with OP50 (either with 0.1% DMSO as control or MK-8722 at the indicated concentration). *C. elegans* were transferred to fresh plates containing vehicle or MK-8722 on days 5 and 10. At least 60 animals were examined for each condition and scored every two or three days until all animals were dead. *C. elegans* that ruptured, bagged, burrowed, or crawled off the plates were censored. Contaminated plates were discarded and excluded from the assay. Lifespan assays were conducted at 20°C.

### Automated lifespan assay

Synchronized L4-stage wild type N2 *C. elegans* were plated on FudR plates seeded with OP50 that was inactivated with paraformaldehyde. The plates were placed on Epson V850 or V800 flatbed scanners and images were collected at regular intervals for 35 days. The protocol for image collection and analysis was based on the Automated Lifespan Machine protocol. At least 150 individual animal lifespans were collected per condition.

### Generation of male *C. elegans*

L4 hermaphrodites were placed at 37 °C for 1 hour and 15 minutes. Males were selected in the progeny after 2 days. To generate sufficient males for the mated brood size assay, hermaphrodites were mated with males and the resulting males were used for the brood size measurement assay.

### Measurement of selfed brood size

Individual L4 hermaphrodites were transferred onto NGM plates seeded with OP50 and left to lay eggs for 24 hours at 20°C. *C. elegans* were transferred to fresh plates every day until they stopped laying eggs. After hatching, the viable progeny produced per individual were counted manually under a light microscope.

### Measurement of brood size of individual mating hermaphrodites

Single L4 hermaphrodites were placed on a plate with 3-4 young male worms. Worms were transferred to a fresh plate every day until they stopped producing eggs. Worms that ruptured, bagged, burrowed, or crawled off the plates were censored. Contaminated plates were discarded and excluded from the assay. Worms that did not mate (based on male and hermaphrodite ratios in the progeny) were censored from the assays. Mating plates were kept at 20°C. After hatching, the viable progeny were counted manually under a light microscope.

### Generation of *C. elegans* carrying *csb-1(ok2335)* and *aak-2* null allele

Strong linkage between the *csb-1(ok2335)* and *aak-2* loci prevented the generation of *csb-1 aak-2* double knockout worms via crossing. We thus used CRISPR-Cas methods to introduce an *aak-2* null allele in the *csb-1(ok2335)* mutant background. CRISPR-Cas genome editing and microinjection were performed as described in (Ghanta et al., 2021) with some modifications. Injection mixture [Alt-R-Sp Cas9 Nuclease V3 (0.25 μg/μL), Alt-R-CRISPR-tracrRNA 0.1 μg/μL, crRNA (0.056 μg/μL, 5`-atcaagtcactggatgtcgt-3`), pmyo2:GFP (1 ng/μL)] was mixed in nuclease-free duplex buffer to a final volume of 20 μL. The injection mix was incubated at 37 °C for 15 minutes, spun down and the supernatant was used for microinjection. Each injected worm was transferred to an individual fresh NMG plate seeded with OP50 and incubated at 20°C. Screening was performed for worms expressing the co-injection marker pmyo2:GFP to ensure successful microinjection. DNA samples of knockout candidates were sent for Sanger sequencing using primer 5`-accataagctgccaaattg-3` and deletions that caused frameshift mutations in *aak-2* were considered knockout worms.

### Mice

All animal experiments were approved by the local animal ethics committees and carried out under license from the Kantonales Veterinäramt Zurich and performed in accordance with local and federal rules and regulations. Mouse health status was monitored regularly according to FELASA guidelines. 17 month-old wild-type male and female C57BL/6JRj mice were ordered from Janvier and acclimated to the facility for one month. One week before performing baseline measurements mice were all given a control purified diet. Following baseline measurements, cages were randomized to either control or MK-8722 treatment groups and the respective diets were given. The control purified diet was Research Diets D12450B and the MK-8722 diet was Research Diets D12450B with MK-8722 compounded at 50 parts per million to achieve a dose of approximately 4mg/kg body weight. Mice were maintained under standard housing conditions at 22°C on a 12-hour reverse light/dark cycle at a density of 4-5 mice per cage. Mice were allowed ad libitum access to food and water throughout the study. Group numbers were: male control; n=7, male MK-8722; n=14, female control; n=20, female MK-8722; n=15.

Body weight and food intake was measured weekly. Blood glucose was measured using a handheld glucometer. A drop of blood for the measurement was collected from the tip of the tail following a six-hour fast. Body composition was measured using a non-invasive EchoMRI machine (EchoMRI, TX, USA). Frailty was scored using the mouse clinical frailty index that includes 31 parameters (Whitehead et al., 2014b). Briefly, each mouse received a score of either 0 (no deficit), 0.5 (mild deficit), or 1 (severe deficit) for each item in the index and the final frailty index score was calculated by dividing the sum by 31. Respiratory exchange ratio was measured using a Promethion-H home-cage indirect calorimetry system (Sable Systems). O2 consumption and CO2 production were monitored at 1-minute intervals for a period of six days. Respiratory exchange ratio was calculated as the quotient of the CO2 produced and O2 consumed. The Weir equation was used to calculate total energy expenditure. Only the final three days of measurements were used for analysis to ensure an appropriate acclimation period to the new caging.

After six months on the diet mice were fasted for six hours then euthanized at approximately ZT18 by exsanguination via cardiac puncture under isoflurane followed by rapid tissue collection. Tissues were quickly weighed then snap frozen in liquid nitrogen and stored and −80°C for subsequent analysis. Blood was allowed to clot in Sarstedt brown-top clotting tubes for 5 minutes at room temperature, then placed on ice until centrifugation at 20,000xg for 20 minutes at 4°C. Serum was collected and stored at −80°C. Serum free fatty acids, urea, creatinine, ALT and AST were measured using Roche Cobas assays according to manufacturer protocols.

### RNA sequencing

RNA from female livers was extracted using the Qiagen RNeasy Plus Mini kit according to manufacturer protocol. RNA quantity and quality was confirmed using NanoDrop and Agilent Bioanalyzer assays. cDNA libraries were prepared using the Illumina TruSeq Total RNA protocol according to the manufacturer’s instructions. cDNA libraries were pooled and sequenced on an Illumina NovaSeq6000, generating approximately 20 million 150bp paired-end reads per sample. Reads were aligned to the mouse GRCm38 reference genome using the align function and annotated using the associated GTF reference file and the featureCounts function from the Rsubread package. Differential expression analysis was performed using the EdgeR and Limma packages in R. Gene set enrichment analysis was performed using the ClusterProfiler package in R.

### Metabolomics

Frozen liver and kidney tissue were ground under liquid nitrogen using a Retsch cryomill. Approximately 25 mg of tissue was weighed into a new precooled tube and extracted with 40:40:20 acetonitrile:methanol:water at a ratio of 40 μL solvent per 1 mg of tissue. Samples were vortexed for 10 seconds, incubated on wet ice for 20 minutes then centrifuged at 21,000 x g for 20 minutes. At least 50 μL of supernatant was transferred to LC-MS vials for analysis.

LC-MS analysis was performed using hydrophobic interaction liquid chromatography coupled to a Thermo Exploris 240 quadrupole-orbitrap mass spectrometer operating in dual scan negative/positive mode. Scans covered m/z 60 to 1000 at 1Hz with resolution of 120,000. Liquid chromatography was performed on a Water XBridge BEH Amide column (2.1 mm x 150 mm x 2.5 μm particle size, 130 Å pore size). Solvent A was 20 mM ammonium acetate, 20 mM ammonium hydroxide in 95:5 water:acetonitrile, pH 9.45 and Solvent B was acetonitrile. Gradients were: 0 min, 85% B; 2 min, 85% B, 3 min; 80% B; 5 min, 80% B; 6 min, 75% B, 7 min, 75% B, 8 min, 70% B; 9 min, 70% B, 10 min, 50% B; 12 min, 50% B; 13 min, 25% B; 16 min, 25% B; 18 min, 0% B, 23 min, 0% B, 24 min, 85% B. The flow rate was 150 μL/min and injection volume was 8 μL.

Analysis of the resulting spectra was performed using EL-MAVEN (Agrawal et al., 2019). Peaks were picked against a validated knows list with metabolite identities matched to standards using retention time and MS1 spectra. The resulting peak intensities were exported for statistical analysis.

### Statistical analysis

OASIS (http://sbi.postech.ac.kr/oasis) was used for the statistical analysis of lifespan assays (Yang et al., 2011).

For mouse measurements with multiple timepoints, ANOVAs were performed including terms for time, sex, diet (Control vs MK) and time-by-diet interaction. For mouse measurements with a single timepoint, ANOVA was performed including terms for sex and diet when effects were compared across sex, or a Mann-Whitney U test was performed comparing diet groups when effects were compared within sex. For metabolomics statistics, nested linear models were generated including terms for diet, sex and diet-by-sex interaction. False detection rate was controlled using a Benjamini Hochberg correction. Statistical tests were performed using R (4.5.3).

